# Mechanistic modelling and Bayesian inference elucidates the variable dynamics of double-strand break repair

**DOI:** 10.1101/026070

**Authors:** M. Woods, C.P. Barnes

**Affiliations:** Department of Cell and Developmental Biology, University College London, England

## Abstract

DNA double-strand breaks are lesions that form during metabolism, DNA replication and exposure to mutagens. When a double-strand break occurs one of a number of repair mechanisms is recruited, all of which have differing propensities for mutational events. Despite DNA repair being of crucial importance, the relative contribution of these mechanisms and their regulatory interactions remain to be fully elucidated. Understanding these mutational processes will have a profound impact on our knowledge of genomic instability, with implications across health, disease and evolution. Here we present a new method to model the combined activation of non-homologous end joining, single strand annealing and alternative end joining, following exposure to ionizing radiation. We use Bayesian statistics to integrate eight biological data sets of double-strand break repair curves under varying genetic knockouts and confirm that our model is predictive by re-simulating and comparing to additional data. Analysis of the model suggests that there are at least three disjoint modes of repair, which we assign as fast, slow and intermediate. Our results show that when multiple data sets are combined, the rate for intermediate repair is variable amongst genetic knockouts. Further analysis suggests that the ratio between slow and intermediate repair depends on the presence or absence of DNA-PKcs and Ku70, which implies that non-homologous end joining and alternative end joining are not independent. Finally, we consider the proportion of double-strand breaks within each mechanism as a time series and predict activity as a function of repair rate. We outline how our insights can be directly tested using imaging and sequencing techniques and conclude that there is evidence of variable dynamics in alternative repair pathways. Our approach is an important step towards providing a unifying theoretical framework for the dynamics of DNA repair processes.

## Introduction

Double-strand breaks (DSBs) are lesions in DNA that occur naturally by oxidative stress, DNA replication and exogenous sources [1, 2]. When left unprocessed or during erroneous repair, they cause changes to DNA structure creating mutations and potential genomic instability [3–8]. To repair DSBs, multiple mechanisms have evolved and are known to include non homologous end joining (NHEJ) [7, 9–17], homologous recombination [18] including single strand annealing (SSA) [19, 20], microhomology mediated end joining (MMEJ) [21, 22] and alternative or back-up end joining (AEJ) [23, 24]. The choice of mechanism depends on the structure of the break point, where simple breaks caused by restriction enzymes are different in structure from those caused by ionising radiation (IR) (reviewed in [25,26]). This affects the probability of error prone repair because mutations are mechanism specific and depending on which mechanism is activated a cell might exhibit chromosome translocations [4, 5], small deletions or insertions [6, 7] or recombination leading to loss of heterozygosity [8]. For example, in mouse, error by SSA causes chromosome translocations [4] and in *Saccharomyces cerevisiae*, NHEJ of simple DSBs is associated with small deletions or insertions [7]. *In vivo* studies of DSBs have suggested that in addition to structural activation arising from the break point, cell-cycle dynamics can also play a role in repair mechanism activation (reviewed in [27]). In particular the choice of mechanism is not fixed at the time of damage and cells exhibit a pulse like repair in human U-2 OS cells [28], a behaviour supported by a molecular basis for cell cycle dependence in NHEJ, mediated by Xlf1 phosphorylation [29].

To understand how mutations are distributed in the genome, it is important to uncover the dynamic activation and interplay between different DSB repair mechanisms. This mutual activation is not fully understood, however the individual repair mechanisms and recruitment proteins of NHEJ, SSA and A-EJ have been documented. NHEJ requires little or no homology, is a mechanism of DNA end joining in both unicellular and multicellular organisms [7] and can exhibit fast repair by the binding of DNA dependent protein kinase (DNA-PK) [15]. In vertebrates, NHEJ initiates the recruitment and binding of several proteins. These have been shown to include Ku70, Ku80, DNAPK catalytic subunit (DNA-PKcs), Artemis and Ligase IV in a cell free system [9]. Ku70 and Ku80 are subunits of the protein DNA-PK. Biochemical and genetic data suggest they bind to DNA ends and stimulate the assembly of NHEJ proteins by DNA-PKcs [10, 12]. Repair proceeds by Artemis facilitated overhang processing and end ligation via DNA Ligase IV [13, 14]. Although well studied, new regulating components of NHEJ are still being discovered, for example the protein PAXX [17]. SSA is slower than NHEJ and in yeast works at almost 100% efficiency for homologous regions of at least 400bp [6]. First described in mouse fibroblast cells [19, 20], during SSA two complementary sequences are exposed through a 5′ to 3′ exonuclease end resection and aligned. Remaining overhangs are then cut by an endonuclease and the DNA is reconstructed by DNA polymerase using the homologous sequences as a template. Some of the components that contribute to SSA have been identified in eukaryotes e.g. the complex MRN consisting of Mre11, Rad50 and Nibrin which facilitates DNA end resection [30]. Following resection, replication protein A (RPA) binds to the DNA and when phosphorylated, forms a complex with Rad52 to stimulate DNA annealing [31, 32]. Similarly to NHEJ, following gap repair, SSA is terminated with end ligation by Ligase III [33]. Data of repair kinetics for mutants defective in Rad52 show limited slow repair in comparison to wild type repair curves in gamma irradiated cells in chicken B line cells [34], suggesting that SSA may be active in the repair of DSBs caused by IR. In yeast, it has been suggested that SSA constitutes a major role in the repair of DSBs accounting for three to four times more repairs than gene conversion during M phase [35].

One interesting finding in genetic studies is that when NHEJ is compromised, DSBs are removed by alternative mechanisms, that have adopted various names in the literature, such as MMEJ in yeast [36] and back-up NHEJ (B-NHEJ) in higher eukaryotes [37] that here, we collectively refer to as A-EJ [23, 38], (reviewed in [37, 39, 40]). It is still unclear how A-EJ is regulated or interacts with other processes but there is evidence it is active in the repair of breaks with microhomology of 3-16 nucleotides, reviewed by Decottignies [40]. Thought to act on break points with ends that are not complementary in the absence of NHEJ factors [38], an assortment of PARP-1, 53BP1, Lig3 and 1, Mre11, CtIP and Pol*θ* have been proposed as regulators of A-EJ. PARP-1 is required and competes with Ku for binding to DNA ends through the PARP-1 DNA binding domain [24]. Other proteins are involved in initial binding, where activation of 53BP1 in MMEJ is dependent on Ku70 and independent of DNA-PKcs [22] and CtIP has been associated through the use of microhomology [41]. The proteins required for end joining have been identified as Lig3 and Lig1 in the absence of XRCC1 [42–44]. This pathway has never been observed in single cells and it is unclear how A-EJ is related to other mechanisms. However, targeted RNAi screening for A-EJ has uncovered shared DNA damage response factors with homologous recombination [45]. For an illustration of the three mechanisms (see Fig. 1a).

**Figure 1:**
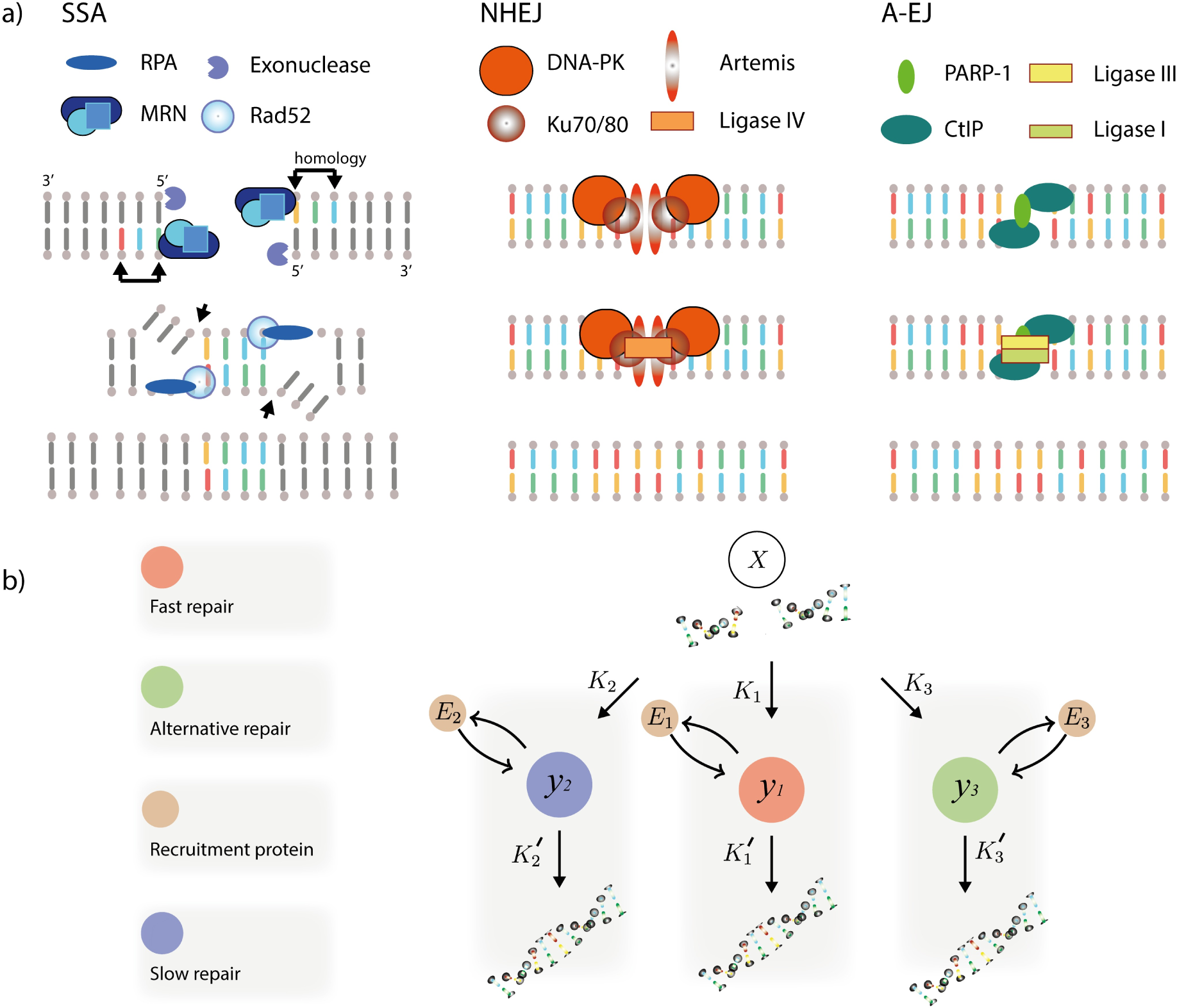
Modelling multiple repair mechanisms. a) Proteins and repair steps contributing to repair during SSA, NHEJ and A-EJ in mammalian cells (illustration). b) The model. Filled circles represent species and arrows represent reactions.

Mathematical models of DSB repair have used biphasic [46], biochemical kinetic [47–52], multiscale [53, 54], and stochastic methods [55]. In a study by Cucinotta *et al*. [48], a set of coupled non-linear ordinary differential equations were developed. The model was based on the law of mass action and stepwise irreversible binding of repair proteins to describe NHEJ rejoining kinetics and the phosphorylation of H2AX by DNA-PKcs. Similar studies have modelled repair kinetics and protein recruitment during SSA [49,51], NHEJ [50] and other mechanisms including NHEJ, HR, SSA and two alternative pathways under a wide range of linear energy transfer (LET) values and heavy ions [52]. These studies have been met with some controversy, for example with the argument that the biphasic model has never succeeded in providing definitive values for the repair components [56]. Recently however, the models have been further developed to model the complexity of a DSB by application to damage induced by ionising radiation of different qualities [57,58]. This can be achieved because the spectrum of DSB-DNA damage can be computed by applying Monte Carlo track structure [59], which is a method that can be used to simulate the passage of charged particles in water, for a review see [60].

Previous biochemical kinetic models have been used to reproduce the experimental data observed. This approach often uses more parameters than are required to describe sequential steps in the repair process. This can cause difficulty in identifying parameter values because multiple parameter value combinations may be able to describe the data well, an issue known as identifiability. Consequently, predictions are not unique, which can be detrimental in the design of a biological experiment. Therefore the creation of models that provide a unique interpretation of repair dynamics is a challenge.

Here we develop a statistical model that can take DSB repair curve data, such as those generated from pulse field gel electrophoresis (PFGE) or comet assays, and infer repair mechanism activation. The method relies on training a simple model against multiple data sets of DSB repair under different genetic knockouts when multiple repair mechanisms are activated. Using the most probable set of parameter values, we can then simulate the model and make predictions on the activation of different rates of repair. Unlike previous modelling approaches, we do not model individual recruitment proteins. Instead we assign parameter values to different rates of repair. This has two benefits. Firstly it provides a method to uncover different rates of repair arising from different repair mechanisms that are implicit in the data. Secondly, it reduces the number of parameters required to describe the system, leading to a more identifiable model. Our approach strikes a balance between a detailed mechanistic description of the biochemical components with a traditional statistical model. This enables insights into the dynamical process underling repair pathways combined with novel and testable predictions. We use this method to integrate the data from eight repair curve assays under genetic knockouts including combinations of Ku70, DNA-PKcs, Rad52 and Rad54. We first infer that there are at least three disjoint dynamical repair mechanisms that explain the combined data and that the dynamics depend on the regulating recruitment proteins. We propose that there are a number of alternative end joining dynamical processes that may or may not share a common genetic pathway. We also demonstrate that our model has predictive power on new data sets, including PARP-1 knockouts, and show that the activation of different repair processes over time depends on the speed of the underlying dynamics.

## Materials and Methods

### Experimental data

The experimental data used in this study are published repair curves generated from methods of pulse field gel electrophoresis, a technique that distributes the DNA according to the length of the fragment. We model the dose equivalent number of DSBs that are obtained from the fraction of DNA released into the gel [61]. Table 1 lists the experimental data that are used for inference. Cells were exposed to X-rays [24] and the number of DSBs within the population recorded over time. The eight data sets are labelled D1-D8. Data D1 is wild type and since the cell cycle phase is unrestricted, we expect all three repair processes to be present. Data D2 and D3 are DNA-PKcs knockouts in G1 and G2 phase, where we expect NHEJ to be compromised but since Ku is present we still expect the recruitment process. Data D4 is a Rad52 knockout where we expect only NHEJ and A-EJ to be present. Data D5 and D6 are Ku knockouts, where we assume the whole of the NHEJ pathway to be compromised and only SSA and A-EJ remain active. Data D7 and D8 are expected to have no repair by PARP-1 mediated A-EJ because both sets were treated with PARP-1 inhibitors. Data D7 comes from *Ku*70^−*/*−^ mouse fibroblasts, where we expect to see no repair by NHEJ as well as a lack of A-EJ due to PARP-1 inhibition [24].

**Table 1:** Table of data sets used for model fitting. The data contains DSB repair kinetics for cells that are irradiated at different doses or split into different phases of the cell cycle, G1 and G2. Data was traced from current literature, or where indicated was provided by G. Iliakis (*). References to the data and cell lines are provided. We chose a combination of mouse embryonic fibroblasts (MEFs) and DT40 cells because DT40 cells remove DSBs from their genome similarly to mammalian cells [63].

| Data | Dose<br>(Gy) | Phase | Mutant, Cell line | Repair |
| --- | --- | --- | --- | --- |
| D1 | 20 | Asynchronous | WT MEFs* [62] | NHEJ, SSA, A-EJ |
| D2 | 20 | G1 | DNA-PKcs<sup>-/-</sup> MEFs* [62] | SSA, A-EJ |
| D3 | 20 | G2 | DNA-PKcs<sup>-/-</sup> MEFs* [62] | SSA, A-EJ |
| D4 | 80 | Asynchronous | Rad52<sup>-/-</sup> DT40* [63] | NHEJ, A-EJ |
| D5 | 80 | Asynchronous | Ku70<sup>-/-</sup>/Rad54<sup>-/-</sup> DT40* [34] | SSA, A-EJ |
| D6 | 54 | Asynchronous | Ku70<sup>-/-</sup> DT40* [34] | SSA, A-EJ |
| D7 | 52 | Asynchronous | Ku70<sup>-/-</sup> + DPQ MEFs [24] | SSA |
| D8 | 32 | Asynchronous | WT + 3'-AB MEFs [24] | NHEJ, SSA |

### Modelling double-strand break repair processes

We assume DSBs caused by ionising radiation (IR) can be repaired by multiple processes with different rates corresponding to NHEJ, SSA and A-EJ (see Fig. 1b). In principle, other mechanisms could be included, such as Rad54 dependent homologous recombination (HR), but since this mechanism is is mostly active in S phase of the cell cycle, and thought to contribute little to IR induced DSBs [62], we chose not to model it explicitly. We model a DNA repair process by a stochastic reaction system, represented by

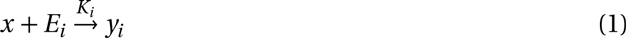

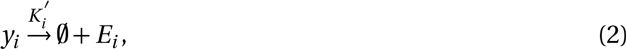

where *x* is the DSB, *E*_*i*_ is the recruitment protein for process *i* (Ku, MRN and PARP-1), and *K*_*i*_, 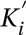 are the parameters for recruitment and subsequent ligation respectively. To model the limited resources available to the cell, we follow the approach of Cucinotta *et al* [48], and assume that the total amount of protein is conserved for each repair mechanism

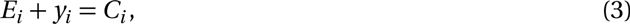

where *C*_*i*_ is the total amount of recruitment protein for each repair mechanism. This structure also captures the experimentally observed fact that mRNA levels for repair processes depend on the radiation dose [64]. The reactions result in a nonlinear coupled stochastic system which is simulated using the Gillespie algorithm [65].

The proportions of DSBs repaired by each mechanism are estimated by calculating the cumulative number of DSBs that enter each individual pathway with the integral

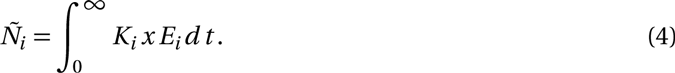

This integral can then be used to calculate the proportion of DSBs repaired by each mechanism.

### Hierarchical modelling of repair across knockout strains

To build a model that can be used to obtain unique predictions, it is advantageous to minimise the number of parameters that describe the system. To do this, we developed a hierarchical model where the individual parameter values, *K*, *K*′, are lognormally distributed with a common mean, *µ*, across all the data sets in which they are included (see Fig. 2a). For data sets in which a repair protein is repressed downstream of the initial protein that binds, we impose an additional hyperparameter. We include this additional hyperparameter because it is not clear if a repair mechanism remains active when individual regulating proteins are repressed.

**Figure 2:**
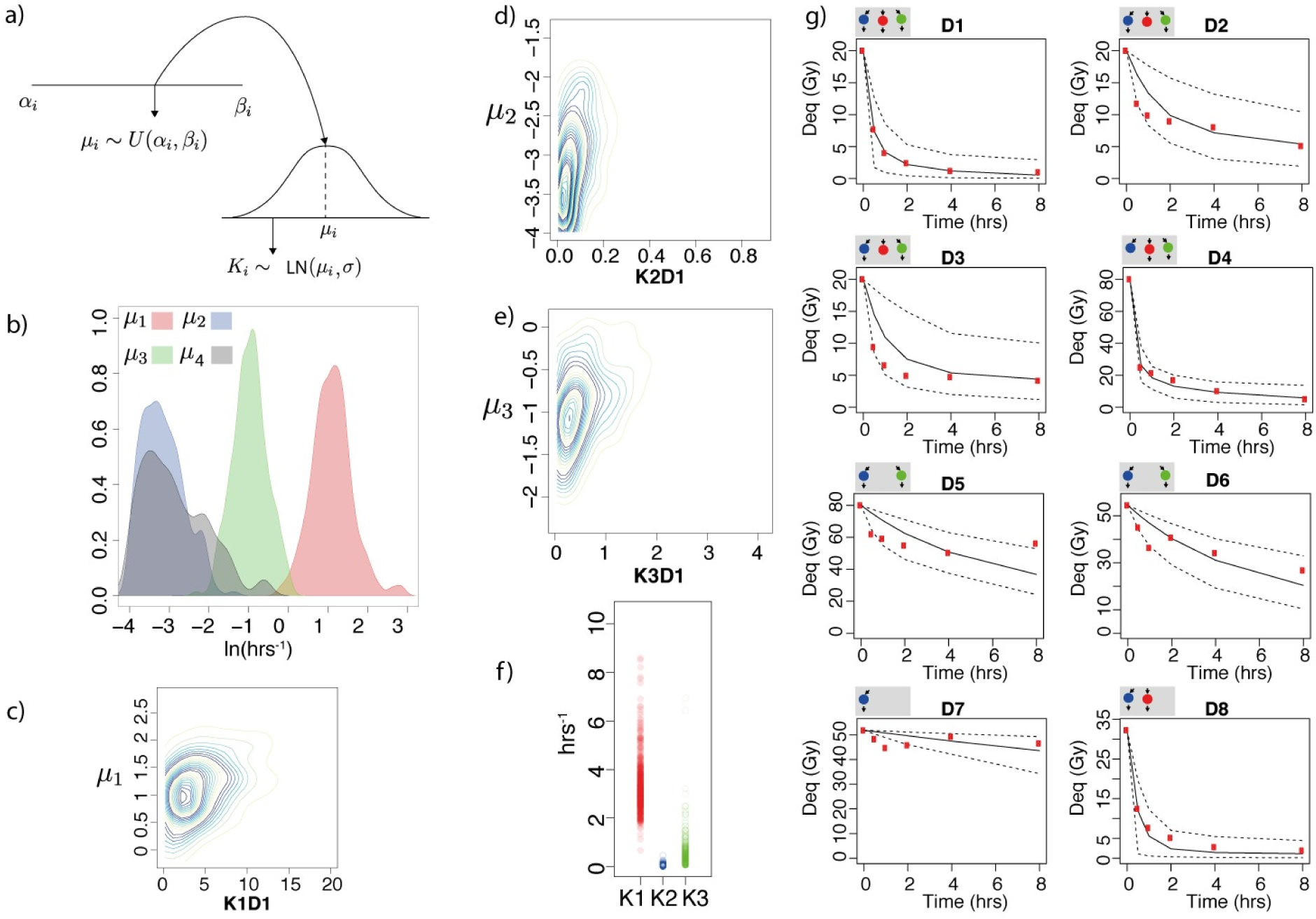
a) Diagram showing the parameter sampling process. Hyperparameters *µ*_*i*_ are drawn from a uniform distribution between *α*_*i*_ and *β*_*i*_. Model parameters *K*_*i*_ are sampled from a lognormal distribution with mean *µ*_*i*_ and variance *σ*^2^. b) Posterior distributions for the hyperparameters *µ*_1−3_ and *µ*_4_. c-e) Posterior analysis for data set *D* 1. Marginal distributions of the parameters *K*_1*D* 1_, *K*_2*D* 1_, *K*_3*D* 1_ against the hyperparameters. f) Posterior distributions of the parameters *K*_1*D* 1_, *K*_2*D* 1_, *K*_3*D* 1_, showing some overlap. g) Time series plots of the experimental data and model fit. Subfigures to the side of each figure represent the active repair mechanisms. The y axis represents the dose equivalent in units of Gray (Gy) (1 DSB = 0.0286 Gy). Red, blue and green represent fast, slow and alternative repair, black solid lines are the median fit and the dashed black lines represent the 0.95 credible regions.

More formally, we wish to obtain 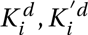 for *i* ∈ {1,2,3} and eight data sets *d* ∈ {1, …,8}. These parameters are constrained by the repair dependent hyperparameters, *γ* = {*µ*_*i*_, *σ*^2^}, where *µ*_*i*_ represents the mean of a lognormal distribution, and *σ*^2^ the variance. The 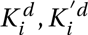 are drawn from the population level distributions, where the joint density can be written

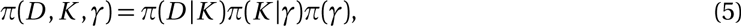

and Bayes rule becomes

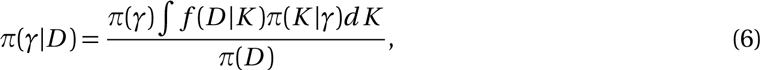

which gives the posterior of the hyperparameters given the data, *D*. The integral indicates that we sum over (marginalise) the *K* values.

### Parameter estimation using approximate Bayesian computation

To perform the inference we use approximate Bayesian computation sequential Monte Carlo (ABC SMC) [66–69]. This method that can be used to fit a model to multiple data sets when the likelihood is unavailable. In the Bayesian framework, we are interested in the posterior distribution 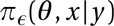, where *θ* is a vector of parameters and *x*|*y* is the simulated data conditioned on the experimental data. To obtain samples from the posterior distribution we must condition on the data *y* and this is done via an indicator function 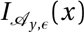. We then have

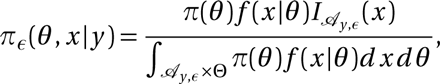

where 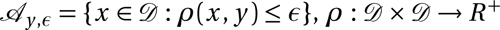 is a distance function comparing the simulated data to the observed data and *π*_*ε*_ is an approximation to the true posterior distribution. This approximation is obtained via a sequential importance sampling algorithm that repetitively samples from the parameter space until *ε* is small, such that the resulting approximate posterior, *π*_*ε*_, is close to the true posterior.

The ABC SMC algorithm is used to calculate an approximation to the target posterior density *π*_*ε*_(*γ*|*D*), where *D* = {*D*_*i*_; *i* = 1,2, &,8}. We can include the hierarchical model by simulating data, *x*\*, using the following scheme:

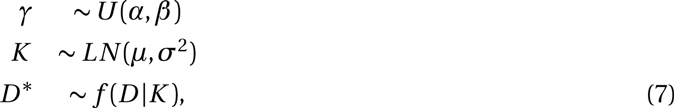

where *f* (*D* |*K*) is the data generating model (the solution to the reaction system represented by Equations 1-3). The *α*, *β* are the lower and upper limits of the uniform prior on the hyperparameters. Note that in the sequential importance sampling step, we perturb only the hyperparameters. For further details on the inference and all prior values see S1 Text.

## Results

### DSB repair requires fast, slow and alternative mechanisms

We fit three different models, comprising one, two and three repair processes respectively (*M*_1_: {*i* = 1}, *M*_2_: {*i* = 1, 2} and *M*_3_: {*i* = 1, 2, 3}), and found that a three process model describes the best fit using an approximation to the Deviance Information Criterion (DIC) [70] and the Akaike information criterion (AIC) [71], based on a surrogate likelihood approach [72] (Fig. B, C in S1 Text). The final model structure is presented in Table 2, (for prior distributions see S1 Text), and a summary of the fitted parameter values is given in Table 3.

**Table 2:**
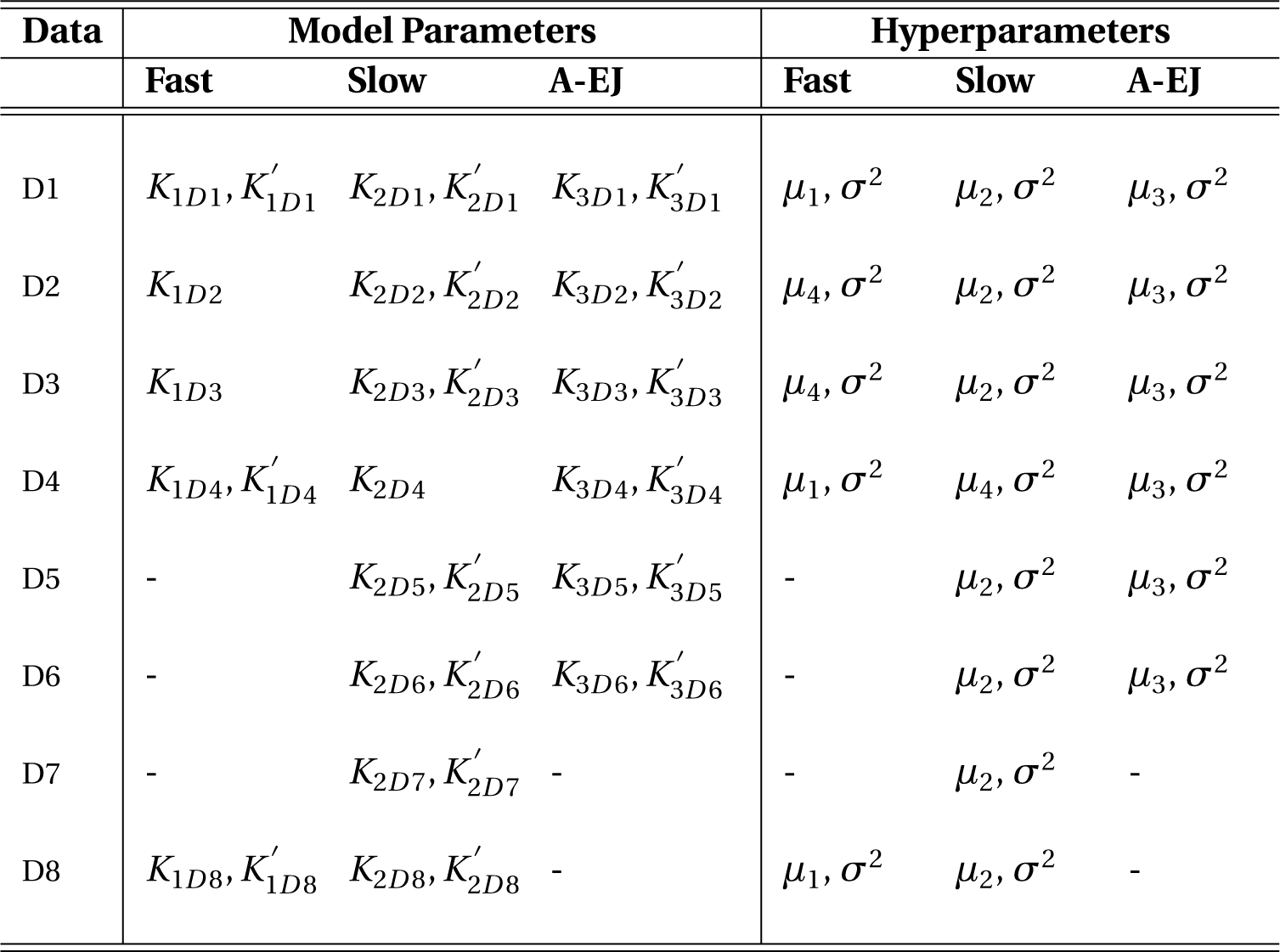
Model parameters with their corresponding hyperparameters used in the hierarchical model. Their values are fitted using ABC SMC. For prior distributions on the hyperparameters, see Text S1.

**Table 3:** Median posterior parameter values with 0.5 credible regions. Values can be compared to initial binding rate constants from two models in the literature; *k* = 0.5 hrs^−1^ for NHEJ [50] and *k* = 0.057 hrs^−1^ for SSA [51]. Half times of DSBs can be calculated using *t*_1/2_ = log_*e*_/*K*. † Cases in which a protein proceeding the first complex to bind is inhibited. In this model we assume there is initial binding but no repair.

| Data | Model Parameters | | | Posterior median ($q_{0.25}, q_{0.75}$) (hrs<sup>-1</sup>) | | |
| --- | --- | --- | --- | --- | --- | --- |
|  | Fast | Slow | A-EJ | Fast | Slow | A-EJ |
| D1 | $K_{1D1}$ | $K_{2D1}$ | $K_{3D1}$ | 3.06 (1.9,5.36) | 0.04 (0.022,0.078) | 0.34 (0.19,0.62) |
| D2 | $K_{1D2}$ | $K_{2D2}$ | $K_{3D2}$ | 0.05 (0.023,0.094) † | 0.037 (0.019,0.067) | 0.4 (0.26,0.63) |
| D3 | $K_{1D3}$ | $K_{2D3}$ | $K_{3D3}$ | 0.04 (0.022,0.081) † | 0.038 (0.021,0.08) | 0.5 (0.31,0.84) |
| D4 | $K_{1D4}$ | $K_{2D4}$ | $K_{3D4}$ | 2.24 (1.52,3.56) | 0.056 (0.027,0.11) † | 0.45 (0.26,0.75) |
| D5 | - | $K_{2D5}$ | $K_{3D5}$ | - | 0.066 (0.034,0.13) | 0.22 (0.13,0.42) |
| D6 | - | $K_{2D6}$ | $K_{3D6}$ | - | 0.057 (0.03,0.102) | 0.29 (0.16,0.53) |
| D7 | - | $K_{2D7}$ | - | - | 0.042 (0.022,0.077) | - |
| D8 | $K_{1D8}$ | $K_{2D8}$ | - | 2.34 (1.75,3.16) | 0.048 (0.03,0.092) | - |
| D1 | $K'_{1D1}$ | $K'_{2D1}$ | $K'_{3D1}$ | 2.87 (2.03,4.49) | 0.04 (0.024,0.08) | 0.4 (0.22,0.65) |
| D2 | - | $K'_{2D2}$ | $K'_{3D2}$ | - † | 0.041 (0.021,0.08) | 0.62 (0.4,0.97) |
| D3 | - | $K'_{2D3}$ | $K'_{3D3}$ | - † | 0.041 (0.023,0.08) | 0.91 (0.58,1.33) |
| D4 | $K'_{1D4}$ | - | $K'_{3D4}$ | 4.35 (3.07,6.57) | - † | 0.29 (0.19,0.46) |
| D5 | - | $K'_{2D5}$ | $K'_{3D5}$ | - | 0.025 (0.016,0.046) | 0.16 (0.12,0.31) |
| D6 | - | $K'_{2D6}$ | $K'_{3D6}$ | - | 0.034 (0.02,0.058) | 0.19 (0.15,0.28) |
| D7 | - | $K'_{2D7}$ | - | - | 0.022 (0.015,0.032) | - |
| D8 | $K'_{1D8}$ | $K'_{2D8}$ | - | 3.2 (2.6,4.4) | 0.041 (0.022,0.085) | - |

The fit of the simulation to the data for all eight data sets is shown in Fig. 2g. The fits capture the essential aspects of the repair curves and most points are consistent with the posterior median and credible regions. When we qualitatively compare this fit to that obtained by the one and two process models, *M*_1_ and *M*_2_, we find a poor fit for *M*_2_ in data sets D2-4 and generally large credible regions for *M*_1_ (see Fig. D, E, F in S1 Text). The posterior distributions of the hyperparameters are shown in Fig. 2b. Inspection of the interquartile range of the hyperparameters confirms that a combination of fast, slow and intermediate repair is sufficient to describe the wild type and mutant data, furthermore a two sided Kolmogorov Smirnov test between the posterior distributions for the hyperparameters confirmed that the four distributions were significantly different to one another (*µ*_1_, *µ*_2_ *D* = 1, *µ*_1_, *µ*_3_ *D* = 1, *µ*_2_, *µ*_3_ *D* = 0.998, *µ*_2_, *µ*_4_ *D* = 0.686, *µ*_4_, *µ*_3_ *D* = 0.752, *µ*_4_, *µ*_1_ *D* = 1, all tests *p* < 2.2*e*^−16^). For each data set (D1-D8) the posterior interquartile ranges of the parameters *K*_1_, *K*_2_ and *K*_3_ were recorded (Fig. G in S1 Text). Marginal distributions for the wild type *K*_1_, *K*_2_, *K*_3_ are shown in Fig. 2 c-e). Analysis of the marginal distribution shows that the parameter distributions of *K*_1_, *K*_2_ and *K*_3_ deviate from the hyperparameter distributions, suggesting that although the rates are defined as fast, slow and intermediate, there is variation in activation of the mechanisms among different mutants (Fig. 2c-e). There is some overlap in parameter values *K*_1_, *K*_2_ and *K*_3_ (Fig. 2f) but the interquartile ranges of the parameters *K*_1_, *K*_2_ and *K*_3_ are disjoint, this is also observed in all eight data sets (Fig. G in S1 Text). For all posterior distributions of the parameters and a plot of individual DSBs and their repair in a wild type model, see Fig. H-K in S1 Text. When individual DSBs are tracked in the model, the DSBs are quickly distributed amongst the three pathways and repaired according to the predicted rate. To check that our model parameters were robust to adding additional data, we performed ABC SMC again on nine data sets. This new data set consisted of the eight data sets listed in Table 1 and an additional repair time series of xrs-6 cells deficient in Ku80 inhibited of PARP-1 by DPQ [24], in which cells were deficient in NHEJ and A-EJ. The results of the total number of DSBs repaired and our predictions on the activation of A-EJ using this data set were the same and a comparison between the eight data and nine data posterior showed a similar fit (Figs. L and M in S1 Text). In summary, we conclude that a three process model provides the best fit to the data observed (Fig. B-C in S1 Text) and that for equal prior ranges on the processes the biological data can be explained by one fast, one slow and at least one intermediate rate of repair.

### The number of DSBs repaired by each mechanism depends on regulating recruitment proteins

By re-simulating from our fitted posterior distribution we were able to examine the dynamics of DSB repair across mechanisms and data sets (see Fig. 3). Data sets in which NHEJ is active exhibited a faster repair with the cumulative number of DSBs reaching to within 80% of the total within a period of 2 hours post irradiation. Next, we plotted the number of DSBs entering each repair mechanism as a time series (Fig. 3b,d). The simulated data predicts that fast repair consistently processes most of the DSBs within two hours after radiation (red curves in Fig. 3). Similarly, there were no clear differences amongst the data in the DSB processing by slow repair. Intriguingly, intermediate repair was slower in cells compromised of Ku70 (D5,D6) than those without DNA-PKcs (D2,D3) (green curves, Fig. 3b,d) To calculate the predicted number of DSBs repaired by fast, slow and alternative mechanisms, we computed the integral 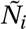. The results are shown in Fig. 4. Data sets for which cells were deficient in regulating components of NHEJ confirmed variation in the numbers of DSBs repaired by intermediate rates. In agreement with the results obtained from the time series plots (Fig. 3b,d) there was a difference in the ratio of slow and intermediate rates between data sets D2,D3 and D5,D6. We also observed an increase in the number of DSBs repaired by A-EJ between G1 and G2 (Figure 4 D2,D3), agreeing with experimental results in the literature [62].

**Figure 3:**
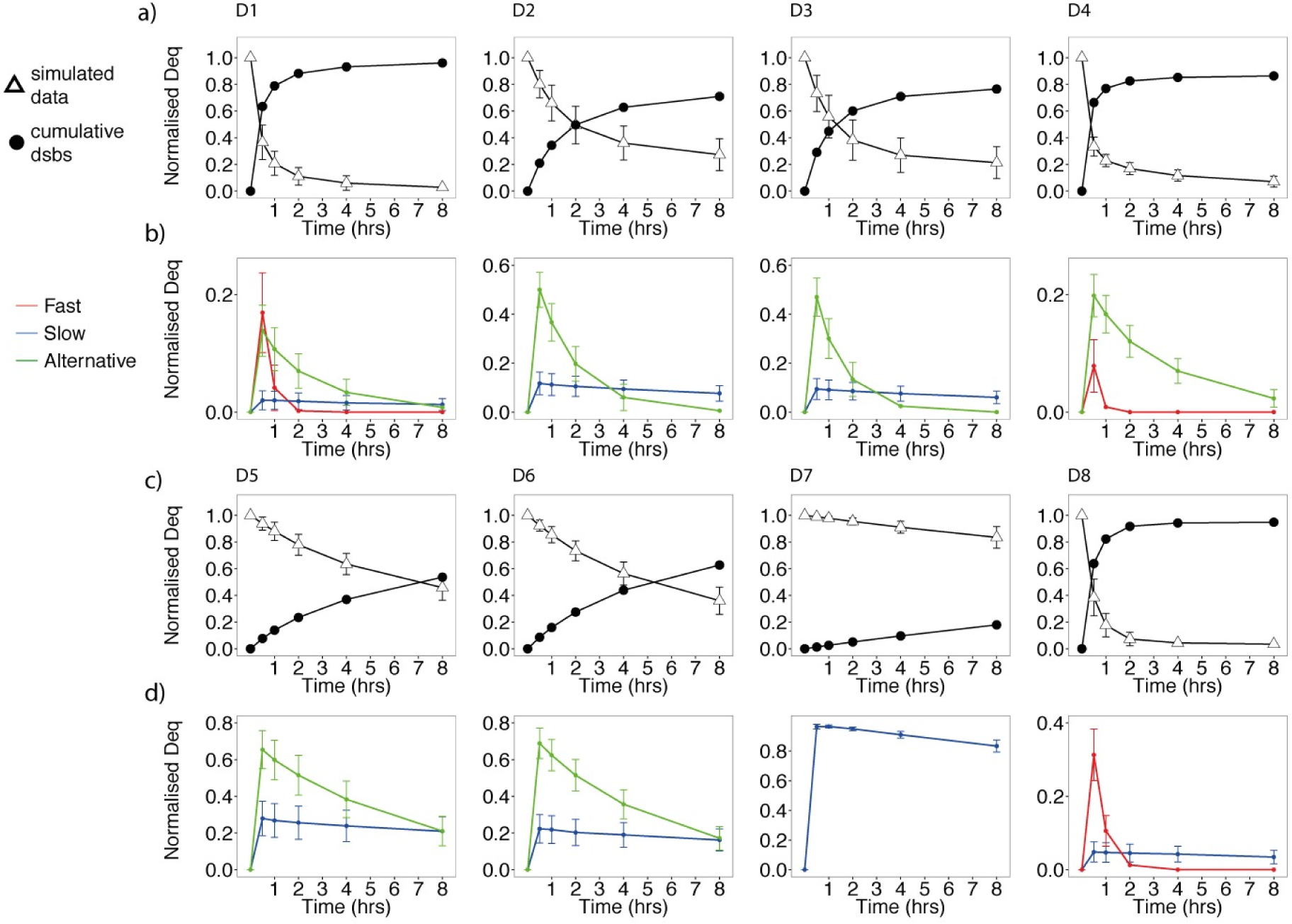
Normalised repair, cumulative repair and the proportion of DSBs within each mechanism as a percentage of the total DSBs for data D1-D8. Plots a) and c) show DSB repair curves simulated from the posterior together with the cumulative number of DSBs repaired. Plots b) and d) show the DSBs entering each repair mechanism. The errorbars represent the 0.5 credible regions (0.25,0.75). Red, blue and green represent fast, slow and alternative repair.

**Figure 4:**
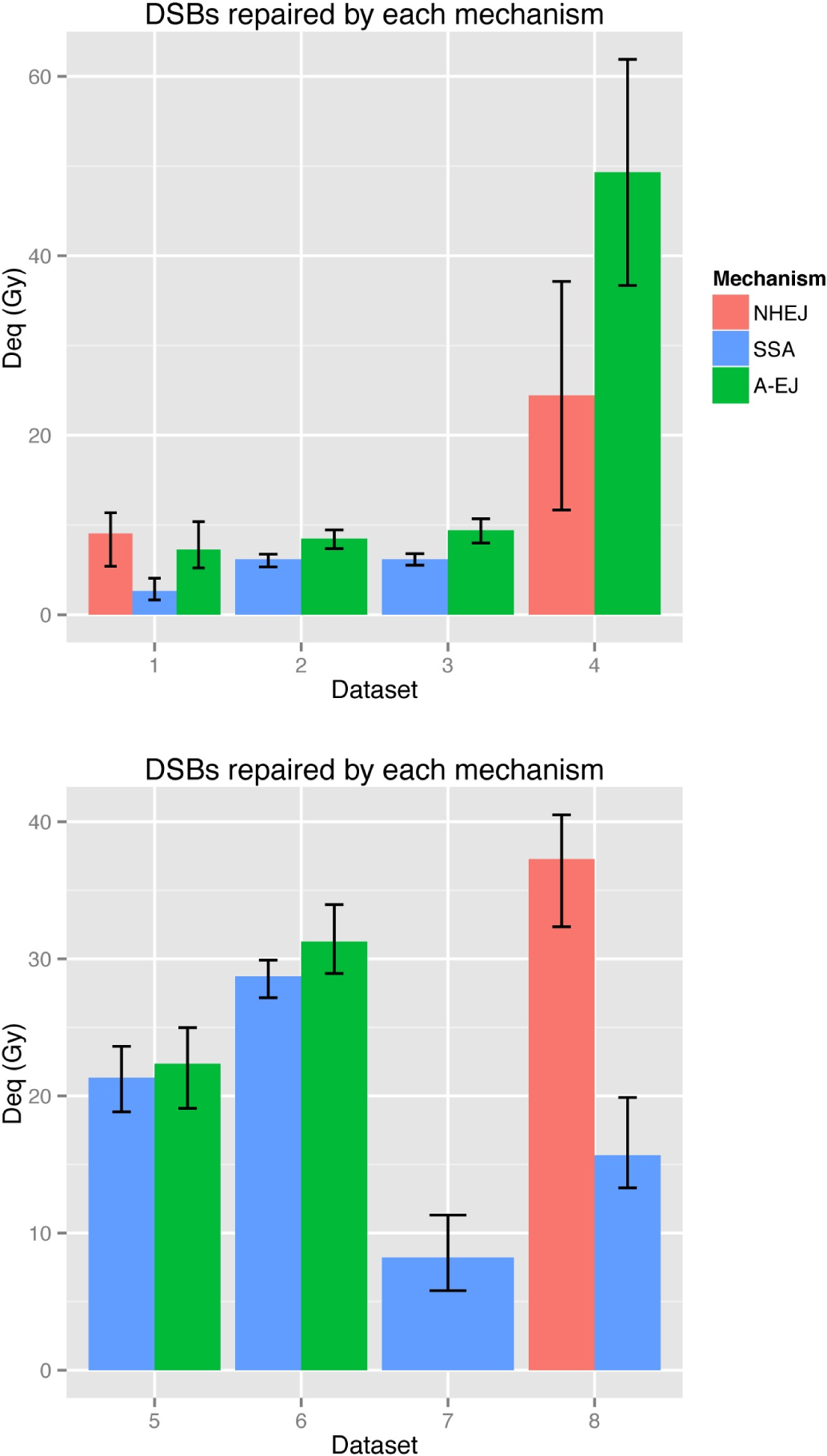
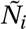, the total number of DSBs repaired by each mechanism for each data set (dose equivalent (Gy)). The errorbars represent the 0.5 credible regions (0.25,0.75). Red, blue and green represent fast, slow and alternative repair.

### Fast and slow kinetics exhibit constrained rate parameters

To test the predictive capability of the model, we re-simulated repair curves using the posterior parameters for data sets D1-D8 and compared the simulated curves to new data sets from the literature (Fig. 5). The model trajectories provide a good fit to wild type data at 40Gy (Fig. 5a, [62]). More impressively, the simulations of cells defective in PARP-1 by application of 3’-AB and Ku80 deficient xrs-6 cells exposed to DPQ defective in PARP-1 and NHEJ show good agreement with the experimental curves (Fig. 5b,c, [24]). This analysis suggests that small ranges of the rate parameters for the fast and slow repair processes, when assigned to NHEJ and SSA, can predict multiple low LET data sets under different experimental conditions. This suggests that they are constrained across experimental and biological conditions. Next we investigated whether the intermediate rates from our model could potentially fit the experimental data when activation of fast and slow rates are inhibited. We re-simulated the model again but this time set the parameters for active NHEJ and SSA to zero across the models for D1-D8. The results are shown in Fig. 5d along with the corresponding experimental data. In wild type data, we see that it is possible for all the DSBs to be removed with the predicted rates of A-EJ, although repair is slower, suggesting that the ability for A-EJ to repair DSBs is not saturated. When Rad52 is inhibited (data D4), because PARP-1 could still compete with MRN we see that it is not possible with the predicted rates for all the DSBs to be repaired in the absence of NHEJ. There was no clear difference in the repair of DNA-PKcs mutants, however when Ku70 is inhibited (data D5, D6) we see that when competition by slow repair is removed the predicted rates of A-EJ, suggest that the activation of A-EJ is not saturated and is perhaps inhibited by competition with MRN.

**Figure 5:**
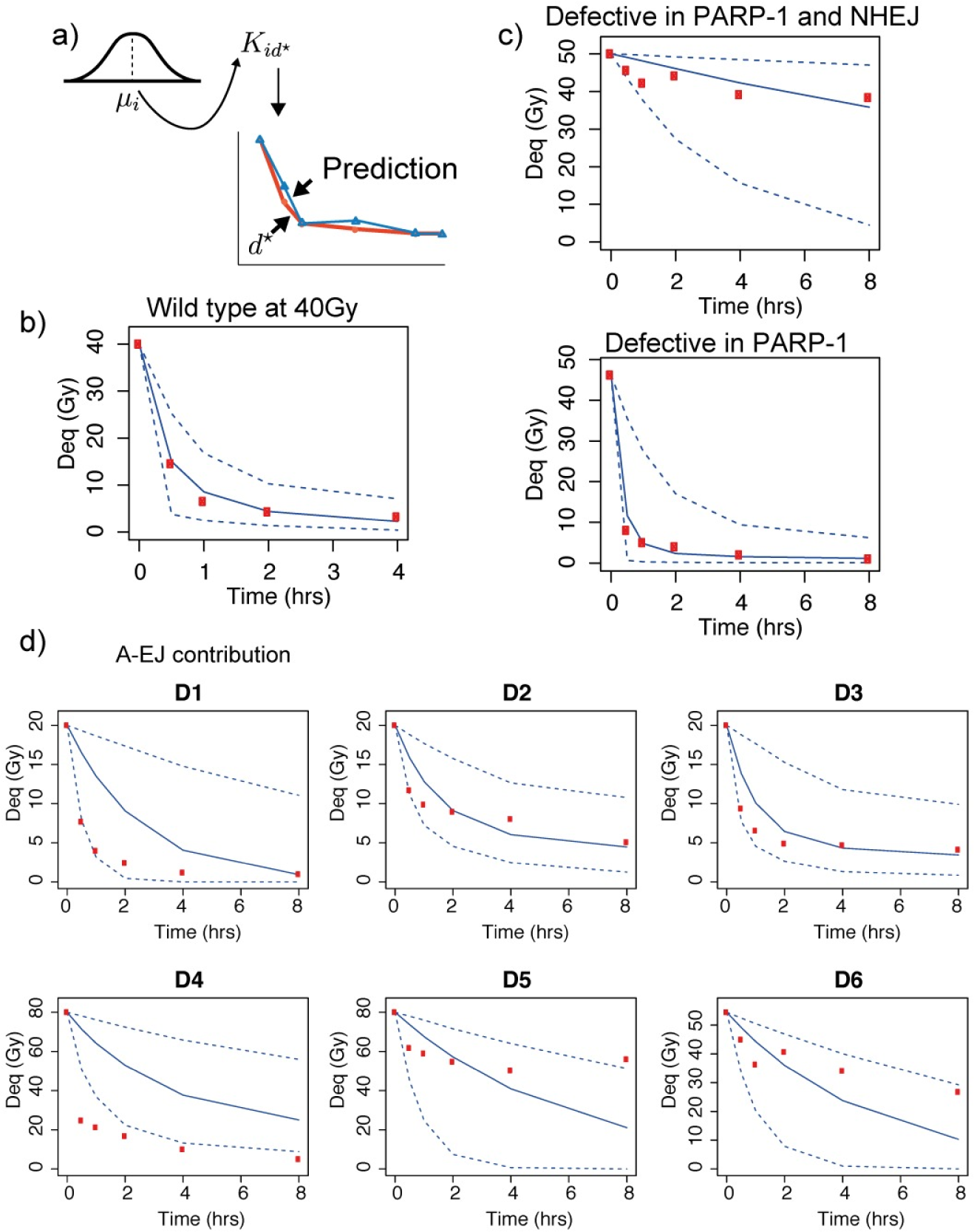
Model predictions. a) Parameters were drawn from the hyperparameter distribution and the code was simulated again under different initial conditions (blue curve) and compared to different data sets (red points) b). Model prediction (blue curve) of wild type repair (red points) from our initial parameter fit (Fig. 2g). c). PARP-1 inhibited repair predictions. The initial parameter fit (blue curves) provided a good fit to the experimental data. d). By setting the parameters of NHEJ and SSA to zero in our predicted posterior, we were able to simulate the expected contribution of A-EJ in our data sets where A-EJ is assumed to be active. Number of DSBs remaining in each of the data sets are shown in the grey shaded region. b-d) Number of DSBs are given in dose equivalent units (Gy). Blue solid lines are the median points, dashed blue lines are the 0.95 credible regions.

### Alternative end joining demonstrates variable dynamics

The time taken for over half the DSBs to be repaired by A-EJ is shown in Fig. 6a. Some repair is fast, occurring within two hours, however, for cells deficient in Ku70, A-EJ adopts a slower repair with half maximum achieved at eight hours. The activation of A-EJ across the data sets is represented by the interquartile ranges of the posterior distributions for *K*_3_ and 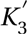 (Fig. 6b). The rate for ligation, 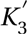, is low in data sets *D*5 and *D*6, suggesting that intermediate mechanisms are less active in the absence of Ku70. The rate is highest in G2 when DNA-PKcs is inhibited. These data suggest that A-EJ adopts a slow or fast repair and that the speed of repair depends on the presence or absence of DNA-PKcs and Ku70, because inhibition of Rad52 had little effect on the time until half-maximum. There are two ways in which this difference between Ku70 and DNA-PKcs mutants can be interpreted. The first is that when Ku70 is inhibited, two alternative mechanisms are activated, one that is fast and one that is slow. The other interpretation is that A-EJ is one repair mechanism that repairs at a slower rate when Ku70 is inhibited. We also modelled the complete inhibition of A-EJ by active NHEJ (A-EJ removed from data sets D1 and D4) and found that the model still captures all the observed dynamics, albeit with the rate of fast repair slightly increased in data set D4 where there is inhibition of Rad52 (see Fig. N in S1 Text).

**Figure 6:**
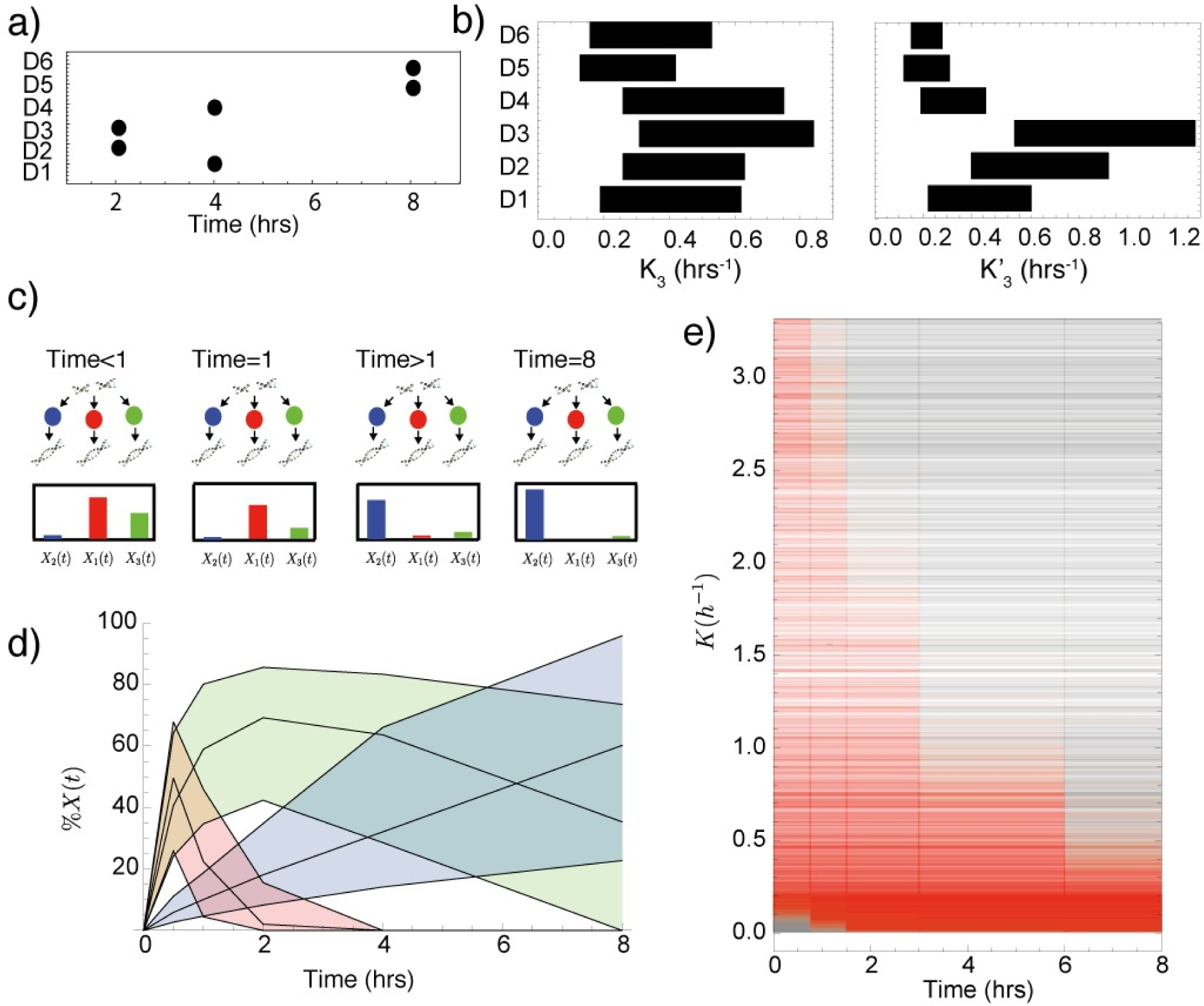
a). The time in which a repair curve reaches below half maximal value for each data set in which A-EJ is assumed to be active. The slowest mode of repair occurred in data sets 5 and 6, where Ku70 is inactive. b). Rectangle plot of the interquartile ranges of *K*_3_ and 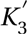 for all data sets where A-EJ is assumed to be active. c). Illustration, showing a typical distribution of the DSBs that remain to be repaired over time. For times < 1 hour a large proportion of DSBs are being repaired by fast NHEJ and faster A-EJ mechanisms, whereas at later times, the majority of DSBs reside in slower HR mechanisms. d). Time series showing the percentage of remaining DSBs in each repair pathway for the wild type data D1. e). Plot showing the time at which each repair mechanism is greater than 30% active for different parameter values. Grey indicates that the mechanism is less than 30% active and red indicates the mechanism is greater than 30% active.

### Total activation time of repair mechanisms depends on the rate of repair

Inspection of the time series data (Fig. 3b,d) suggests that at time *t* = 0.5hrs, the majority of DSBs that are being processed are within a fast mode of repair. Fig. 6c illustrates a typical distribution of DSBs over each mechanism at different points in time. At time *t* = 0, the cells are exposed to a single dose of ionising radiation. Quickly, for example at time *t* < 1hrs, fast and possibly faster alternative repair mechanisms dominate the DSB processing. Later, after all DSBs processed by the faster mechanisms have been repaired, the remaining DSBs fall within the category of breaks that require processing by slower mechanisms. This change in the activity of repair mechanisms could potentially be investigated by recording changes in the level of recruitment proteins or gene expression as time series. To quantify this change in our simulated data, we plotted the percentage of DSBs that remain in active repair mechanisms over time for the wild type data (see Fig. 6d). By inspection, we can see that at 0.5 hours after irradiation most DSBs reside in the fast mechanisms. At a time of *t* = 8hrs, the percentage approches zero for fast repair and the majority of DSBs are found within the slow repair pathways. The variation in Fig. 6d) is shown with the 25th and 75th percentiles and this is due to the variation in repair rate *K*_*i*_, 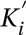 for each mechanism. Ultimately, it is the values of the parameters *K*_*i*_, 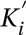 that determine the rate of repair, so to confirm if the dynamics presented in Fig. 6c are representative of the whole data set, we considered all time series for parameters *K*_*i*_, a total of 9000 simulations. For each parameter at every time point, we assigned a value of 1 if the corresponding mechanism for the parameter contained over 30% of the total DSBs being processed at that time point and a value of 0 if it contained less than 30%. The results are shown in Fig. 6e, where for each parameter *K*_*i*_, a red line indicates the times at which the mechanism with rate *K*_*i*_ is greater than 30% active. There is a clear trend showing that the percentage of total activation decreases in time with an increase in repair rate *K*. In other words, the model predicts the times at which different repair processes are likely to express regulating components. When repair is extremely slow the repair mechanism never reaches 30 % of the current DSBs. In summary, these results predict that if a cell experiences a sudden creation of DSBs, then gene expression for slower repair mechanisms will be maintained for longer than those required for faster repair mechanisms such as NHEJ, a result that has been implied for NHEJ and HR (Fig. 3 in [28]).

## Discussion

In this study, we presented a new hierarchical model of DSB repair and applied Bayesian inference to infer the number of active repair processes and their dynamical behaviour from experimental PFGE data. Because the model assumptions are simple and exclude the full mechanistic details of the biological processes, we are able to form an identifiable model and provide unique insights on the difference in dynamics under multiple knock-out cell lines.

We have identified four major insights, each of which can be further tested experimentally. The first insight is that the data is explained by at least three independent mechanisms. Our results suggest that there are multiple dynamic regimes for the intermediate process. For example a mechanism faster than Rad52 dependent HR is required to fit the experimental data to the model in data sets D2 and D3 (knockout of DNA-PKcs). Another interesting finding is that intermediate repair is increased in G2 phase of the cell cycle. If we assume that intermediate repair corresponds to alternative end joining, then this is in agreement with experimental results in the literature, supporting the existing biological evidence of the role of A-EJ in DSB repair [62]. This agrees with genetic studies that suggest two forms of alternative end joining depending on the presence of microhomology [40].

Our second insight is that the speed of A-EJ depends on the presence of regulating components in NHEJ and SSA, and in particular we observe a slower rate when Ku70 is inhibited and a faster rate when DNA-PKcs is inhibited (data sets D5-D6 and D2-D3 respectively, Fig. 3). As reported by experimental analysis, Ku deficient cells do not produce NHEJ products due to excessive degradation or inhibition of end joining [11], and our model suggests that a slower A-EJ is active in these cells (data sets D5-D6). In addition, inhibition of DNA-PKcs does not activate repair by PARP-1 mediated A-EJ [24] and leads to elevated levels of resection and more HR [73]. Together with our model, this suggests that an alternative mechanism that is faster than PARP-1 mediated A-EJ could be activated when DNA-PKcs is inhibited (data sets D2-D3). These results could be tested by examining DSB repair in single cells with and without inhibitors using time-lapse microscopy and existing markers such as fluorescently tagged 53BP1, a protein that co-localises with DSBs, and fluorescent tagged PARP1, a candidate protein for A-EJ [28, 74].

The third insight that is generated from our analysis is the prediction of the total number of DSBs repaired by each mechanism. We can use this to estimate the proportion of different mutations following DSB repair in wildtype and mutant cells. Some cancers are deficient in at least one repair mechanism and in these cases, alternative mechanisms of repair have been observed to compensate [75]. One example is the increase in chromosomal aberrations observed in cells compromised of NHEJ by loss of Ku80 [3]. Recently, mutations specific to alternative mechanisms have been identified, where next generation sequencing has revealed sequence specific chromosome translocations following A-EJ at dysfunctional telomeres [5]. In addition, A-EJ is error prone, giving rise to chromosome translocations, of which there are more when NHEJ is inactive, suggesting it’s role as a back up mechanism in eukaryotes [44]. If we know how many DSBs are likely to be repaired by each mechanism, this information will be important in predicting the numbers and types of mutations that we expect to observe. Potentially, a better understanding of the interplay between DSB repair mechanisms could be applied to design potential synthetic lethal therapeutics in cancer [76].

The fourth insight is that the gene expression profile of the proteins within different DSB repair mechanisms should change over time, with slower repair mechanisms still remaining active many hours after the initial dose of radiation. Pulse-like behaviour has been recorded in the repair of DSBs in human cells [28] and we suggest that this prediction could be further investigated using microarrays or sequencing. In fact, the expression of repair pathway genes has recently been used to diagnose the prognosis of carcinomas [77]. Currently the genes involved in the different repair pathways - and how much they are shared - remains to be fully elucidated. Our model could be used to predict the times at which different repair pathways dominate and provide a theoretical model for the interpretation of time-course gene expression results.

The model will require further development before it can be applied to more general problems in the DNA damage response. With additional data it will be possible to extend the model and include additional terms such as explicit repressive cross-talk interactions. It may be possible that inhibition of a certain protein may not completely ablate the function of a repair pathway. While our two-step model somewhat accounts for this, the actual pathway contains many different proteins and more complex effects could arise. Additionally, the PFGE assay provides a population average of the total repair; in order to obtain information on relatively small numbers of double strand breaks, together with estimates of cellular heterogeneity and stochasticity, data obtained by methods such as live cell imaging are required. Another current limitation arises from the heterogeneity of the DNA damage spectrum, where complex breaks can account for 30-40% of the population [78]. Previous methods of Monte Carlo track structure have been able to predict the number of single strand breaks, simple and complex DSBs that are created [59]. By identifying proteins responsible for the processing of complex DSBs and analysing knockout repair curves, it will be possible to gain understanding on the repair of heterogenous breaks, and incorporate these into our model. Despite our demonstration that the model is predictive across multiple doses in the low LET regime, we modelled repair following a variety of radiation doses, and we can’t rule out that complexity of DSBs further alters the dynamics of repair. Previous studies have also modelled the formation of foci by summing the DSB enzymes involved in the phosphorylation of H2AX [48,52] and using this approach, our model could be developed to take into account lower doses of radiation or the dynamics of *γ*-H2AX foci formation. Related complicating factors are clustered DSBs [79] and DSBs residing in heterochromatic regions, which have been shown to require Artemis for repair [80].

From our simple assumptions we have built a predictive model and generated *in silico* data that was used to produce a number of unique insights that can be tested experimentally. Mathematical modelling not only facilitates the analysis of disparate data sets but also enforces the explicit formalisation of the underlying assumptions of our hypotheses. Our framework is another step towards a theoretical understanding of the dynamics of DNA repair pathways. The DNA damage response is comprised of a large number of extremely complex interacting biological processes. As the collection of larger and more heterogeneous data sets increases, we anticipate that mathematical modelling approaches will be absolutely essential for the reverse engineering and understanding of these complex biological processes.

## Funding

This work was supported by a Wellcome Trust Research Career Development Fellowship [WT097319/Z/11/Z].

## Acknowledgements

We thank Geraint Thomas, David Hall and all members of the CSSB group for helpful discussions, Gyorgy Szabadkai and Panos Zalmas for critical review of the manuscript, and M. Berger at NVIDIA Corporation and Fabrice Ducluzeau at University College London for hardware support. The authors would like to acknowledge that the work presented here made use of the Emerald High Performance Computing facility made available by the Centre for Innovation. The Centre is formed by the universities of Oxford, Southampton, Bristol, and University College London in partnership with the STFC Rutherford-Appleton Laboratory.

